# Climate influences the response of community functional traits to local conditions

**DOI:** 10.1101/2020.10.30.362525

**Authors:** Laura Melissa Guzman, M. Kurtis Trzcinski, Ignacio M. Barberis, Régis Céréghino, Diane S. Srivastava, Benjamin Gilbert, Valério D. Pillar, Paula M. de Omena, A. Andrew M. MacDonald, Bruno Corbara, Céline Leroy, Fabiola Ospina Bautista, Gustavo Q. Romero, Pavel Kratina, Vanderlei J. Debastiani, Ana Z. Gonçalves, Nicholas A.C. Marino, Vinicius F. Farjalla, Barbara A. Richardson, Michael J. Richardson, Olivier Dézerald, Gustavo C.O. Piccoli, Merlijn Jocqué, Guillermo Montero

**Author notes:** Authors contributed equally (Laura Melissa Guzman, M. Kurtis Trzcinski). Correspondence: Laura Melissa Guzman.

## Abstract

Functional traits determine an organism’s performance in a given environment and as such determine which organisms will be found where. Species respond to local conditions, but also to larger scale gradients, such as climate. Trait ecology links these responses of species to community composition and species distributions. Yet, we often do not know which environmental gradients are most important in determining community trait composition at either local or biogeographical scales, or their interaction. Here we quantify the relative contribution of local and climatic conditions to the structure and composition of functional traits found within bromeliad invertebrate communities. We conclude that climate explains more variation in invertebrate trait composition within bromeliads than does local conditions. Importantly, climate mediated the response of traits to local conditions; for example, invertebrates with benthic life-history traits increased with bromeliad water volume only under certain precipitation regimes. Our ability to detect this and other patterns hinged on the compilation of multiple fine-grained datasets, allowing us to contrast the effect of climate vs. local conditions. We suggest that, in addition to sampling communities at local scales, we need to aggregate studies that span large ranges in climate variation in order to fully understand trait filtering at local, regional and global scales.

## Introduction

Ecologists are reformulating long-held perspectives on biodiversity using functional traits. Since organisms interact with their environment through their traits, patterns in species distribution should be a direct function of their traits. Traits directly affect community assembly and species interactions, such that any snapshot of a community is the result of i) physiological tolerances (Winemiller et al. 2015, Pianka et al. 2017), ii) species interactions (Chesson 2000, Chase and Leibold 2004, Estes et al. 2011), Hi) dispersal and priority effects (MacArthur and Wilson 1967, Diamond 1975, Hanski 1999), iv) demographic stochasticity (Lande et al. 2003, Hubbell 2011), and v) phylogenetic constraints (Vellend 2016, Pianka et al. 2017). Most of these mechanisms are, to some degree, driven by functional traits (mechanisms i-iii) or determine the distribution of traits (mechanism v), and their prevalence and diversity modulate the relationship between biodiversity and ecosystem function rather than species *per se* (Schmitz et al. 2015). Similarly, the redundancy of functional traits in a community can maintain ecosystem function and stability in the face of environmental change and species loss (Loreau et al. 2001). By shifting our focus away from taxonomic diversity to the diversity of functional traits within a community or metacommunity, we strengthen our ability to detect the mechanisms that underlie observed patterns in species distribution and biodiversity (McGill et al. 2006). In particular, functional traits have been associated with broad biogeographic patterns, such as the latitudinal gradient in biodiversity, leading to new insights into the processes and causes of global biodiversity (Chave et al. 2009, Ricklefs 2012, Lamanna et al. 2014).

This trait-based paradigm recasts community ecology’s central question about species diversity and coexistence as: which processes determine the functional trait composition of ecological communities? Spatial scale is implicit in this question, as different processes are expected to act at different scales (Levin 1992, Chave 2013). For example, species interactions are expected to be strongest at small spatial scales, whereas environmental filtering often occurs at spatial scales large enough for environmental gradients to exceed physiological thresholds (Kraft and Ackerly 2010). Finally, processes like dispersal limitation and biogeographical constraints to the species pool often operate at the largest spatial scales (Ricklefs and Schluter 1993). The grand challenge of integrating these multiple scale processes on traits can only be resolved by a synthesis of traditional small-scale ecology with maciroecology and biogeography (Violle et al. 2014).

Environmental conditions could have cross-scale effects on functional traits in several ways. First, local factors (e.g. resource availability) could scale up to affect the geographic distribution of functional traits, or, local conditions such as microclimate may compensate apparently limiting climatic conditions. Second, climatic and biogeographic factors could constrain the local distribution of functional traits and thereby impact local processes. A practical difficulty of including large scale environmental factors or biogeography into syntheses of local scale studies is that some variables will be spatially pseudo-replicated and others will not. That is, several field studies that fall within a single climatic zone or geographic region may not represent independent measures, yet a large number of sampling units are needed to detect local effects on community functional trait composition. These issues can be partially resolved by a spatially structured hierarchical analysis.

The theory and motivation for our study could be applied to almost any ecological community. Yet, the data required to test this theory requires extensive information on species composition, functional traits, local conditions and climate for multiple georeferenced occurrences of a defined community across a broad geographic range; such data are rarely available for multitrophic communities. Here we use the aquatic macroinvertebrates in tank bromeliads as a model system to understand community structure. The invertebrate communities in tank bromeliads have proven to be useful systems for testing ecological theory, as they are easily manipulated and censused and are naturally highly replicated. Previous work has related ecosystem functions, trophic structure and system resilience to a variety of invertebrate functional traits (Dézerald et al. 2015, 2018, de Omena et al. 2019). To date, researchers of this system have primarily focused on local-scale explanations for community composition and ecosystem function, detailing how bromeliad size (volume of aquatic habitat), detrital inputs, and canopy cover affect community composition (Petermann et al. 2015, Kratina et al. 2017). However, much of the geographic variation in invertebrate composition remains unexplained, and consequently begs for more explicit incorporation of broadscale variables like climate. For example, extreme rainfall events lead to an inversion of the trophic pyramid of macroinvertebrates in bromeliads across seven study sites broadly distributed across the neotropics (Romero et al. 2020)).

Here, we used fine-grain data on the functional traits of aquatic macroinvertebrates sampled in more than 1600 tank bromeliads across 18 climate zones throughout the Neotropics to partition the separate and combined effects of local conditions and climate on community trait composition. Based on previous studies of bromeliad macroinvertebrate ecosystems, we expect that both local conditions and climatic gradients affect trait composition (Céréghino et al. 2011, 2018, Dézerald et al. 2015). We tested three hypotheses of how trait variation could be partitioned between local and biogeographical environmental gradients: (i) Variation in trait composition is primarily explained by variation in the environment at the biogeographic scale. This would occur when climatic factors, which vary only at biogeographic scales, determine the cost and benefits of different functional traits. For example, temperature determines development rates (Damos and Savopoulou-Soultani 2012) and predation pressure (Romero et al. 2018), while precipitation determines the mortality rates due to desiccation (Amundrud and Srivastava 2015). (ii) Variation in trait composition occurs primarily along regional environmental gradients, or the interaction between local conditions and the climate at a given region. This would occur when heterogeneity in local conditions determines life history traits. For example, traits may vary as a response to the availability of resources within a bromeliad (Srivastava et al. 2008) and the avoidance of negative species interactions (Hammill et al. 2015). (H) Neither local nor regional conditions determine trait composition. This would occur at the regional scale if strong biogeographic constraints dampen any effect climate may have on traits. For example, geographic dispersal barriers on particular clades (Amundrud et al. 2018), coupled with deep phylogenetic signal on some traits override the effects of climatic conditions. Similarly, at a local scale, stochasticity in colonization coupled with noise added by a generalist predator can overwhelm any signal caused by environmental gradients (Srivastava and Bell 2009). We used hierarchical analyses to distinguish between these three hypotheses by testing the effects of local conditions, climate, and their interaction on the community functional traits of bromeliad invertebrates at local and bioclimatic scales.

## Methods

### Invertebrate traits

We used as functional trait values the species scores on four main axes of invertebrate trait variation identified by Cereghino et al. (2018), which represent life-history strategies along trophic, habitat, defence and life-history niche axes describing ca. 852 aquatic invertebrate taxa occurring in Neotropical tank bromeliads, mostly identified to species or morphospecies (hereafter “species”). For completeness, in the following we explain the method used by Cereghino et al. (2018) to identify such a synthetic traits: Each of these species was characterized by 12 nominal traits: maximum body size, aquatic developmental stage, reproduction mode, dispersal mode, resistance forms, respiration mode, locomotion mode, food, feeding group, cohort production interval, morphological defence, and body form. Each nominal trait had a number of modalities, or states. For example, the states for the trait “feeding group” were “deposit feeder”, “shredder”, “scraper”, “filter-feeder”, “piercer” and “predator”. A full description of traits and states can be found in Céréghino et al. (2018). Information on these traits was structured using a fuzzy-coding technique (Chevenet et al. 1994): scores ranged from “0” indicating “no affinity”, to “3” indicating “high affinity” of the species for a given trait state. Scores were based on observations of specimens (Dézerald et al. 2017), on the scientific literature (e.g. Merritt and Cummins 1996, Céréghino et al. 2011) and expert opinion (see Céréghino et al. 2018 for list of traits, modalities, and their definitions) the species x trait matrix is available at https://doi.org/10.5281/zenodta.1200194. Principal Component Analysis (PCA) was used to reduce trait dimensionality to significant axes of trait variation. The rank-transformed [species x trait] matrix was used to compute Spearman’s rank correlations between trait modalities, which then underwent PCA with bootstrap resampling (Pillar, 1999). This procedure allowed us to test ordination stability and to interpret the significant ordination axes in light of correlations with trait states. We identified a robust set of four orthogonal and important axes of trait variation, namely trophic position, habitat use, morphological defence, and life cycle complexity (Céréghino et al. 2018). The species scores for these four PCA axes (available at https://doi.org/10.5281/zenodm.1200194) thus represented continuous trait values, or synthetic traits, which we then used in analyses of the processes underlying functional diversity across different spatial scales in relation to environmental factors (this study).

### Site sampling

The data for the present study consists of the abundance of macroinvertebrates found within 1436 bromeliads in 18 field sites in 8 countries (IBromeliad Working Group BWG database, Table A1). While the Bromeliad Working Group has sampled more countries and field sites, we only use the sites where more than 15 bromeliads were sampled. Some of these field sites were visited in multiple years (Table A1), while others were visited only once. While we acknowledge there are temporal trends in community abundance and composition, all of our sampling units (each sampled bromeliad) are a snap-shot of the community structure, so we made the simplifying assumption to treat all sampling units within a site the same, regardless of the year in which they were collected. Examining temporal trends in trait composition, would be an interesting follow up study.

The full suite of 19 bioclimatic variables was extracted from WorldClim using the latitude and longitude of each sampled site (Fick and Hijmans 2017). Some of our sampling locations were within 1 km^2^ of one another. As such, these locations had the same climatic conditions since this is the smallest resolution in WorldClim. We decided to group the data collected from these locations, instead of attempting to downscale the climatic conditions to a smaller resolution. After grouping the data from these locations, we ended up with 18 sites, which we refer to as bioclimatic zones.

### Bromeliad sampling

The following bromeliad genera were sampled across all sites: *Neoregelia, Quesnelia, Tillandsia, Guzmania, Vriesea, Aechmea* and *Catopsis.* Each tank bromeliad was exhaustively sampled either by dissecting the rosette, or by pipetting out all of its contents. The bromeliad macroinvertebrate community is defined as all the aquatic invertebrates found by the naked eye (> 0.5 mm) within a single plant. A total of 637 morpho-species were found.

Three local conditions were collected for each plant: water volume at time of sampling (mL), total amount of detritus (mg), and canopy cover (either open or closed canopy). The data on total amount of detritus and the water volume at the time of sampling were log-transformed before the analyses. These variables are proxies of habitat size and energy inputs, which are key drivers of food web structure (Oertli et al. 2002, Moore et al. 2004)). The total amount of detritus was calculated by adding the amount of small, medium and large detritus. In 25.8% of bromeliads, total detritus was not measured directly, instead we imputed total detritus with allometric relationships using other size categories of detritus, and in a few cases with the number of leaves and the diameter of the pant. In 50.4% of bromeliads, total water volume was not measured directly and instead we imputed total water volume using leaf size, plant species, plant height, plant diameter, and number of leaves. When either of these variables were missing, we used generalized linear models with a Gaussian error distribution to impute missing values. Thus, the dataset was a combination of directly measured and estimated values.

### Spatial scale of environmental variation

To better understand the spatial scale of environmental variation, we partitioned the variation in local environmental conditions of a bromeliad into site and bioclimatic scales using three nested hierarchical models, one for each response. All three models used the same structure of random effects, but different likelihoods according to the environmental variable. The environmental variables partitioned in this way included log detritus (normal likelihood) log water volume (normal) and canopy cover (binomiall). By using nested random effects, we can partition the variation of each local environmental condition by spatial scale and determine at which spatial scale most of the variation is explained. For each environmental variable, we estimated random effects for the field visit (site by year combination) within bioclimatic zone, and bioclimatic zone. We also calculated the correlation between all local conditions both within and across all bioclimatic zones.

### Trait analysis

Since our unit of analysis is the bromeliad invertebrate community, it was necessary to quantify the presence and abundance of the animals and their traits for each bromeliad. To do this, we calculated the community weighted means (CWM) of each synthetic trait for each bromeliad (local scale analysis). Community weighted means (CWM) was given by:

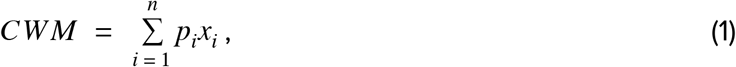

where p_i_ is the proportion of species abundance, x_i_ is the trait value of that species and n is the number of species in that community (Garnier et al. 2004, Swenson 2014). To reduce the effect of highly abundant species on the analysis, we applied a square root transformation of the proportional abundance of invertebrates within a bromeliad (i.e. a Hellinger transformation). This transformation was necessary since we used abundance and not biomass, and the most abundant species were orders of magnitude (up to 100s of individuals) more abundant than the least abundant species (1 or 2 individuals) (Legendre and Gallagher 2001). The square root transformation de-emphasizes superabundant species (gave a more equal weight to rare species), and the community weighted means (CWM) allowed us to characterize the relative abundance of traits in the sampled bromeliad.

Community weighted means were obtained using the data version 0.7.7 extracted from the BWG database in July 2017.

To determine the effects of environmental conditions on the CWM computed using the species scores on the four synthetic traits, we used a permutational multivariate analysis of variance - PERMANOVA (vegan R package; Anderson 2001). This method is based on within- and between-group sums of squares computed on pairwise dissimilarities, in this case, of bromeliad communities considering the CWM trait values. However, instead of permuting the site matrix (bromeliads or bioclimatic zones), we adapted the method to permute among the species vectors in the trait matrix and recomputed the CWMs, to reduce the risk of type I error (Peres-Neto and Kembel 2015, Hawkins et al. 2017, Zelený 2018). Since each bioclimatic zone differs in the number of bromeliad communities sampled (Table A1), and sample size may bias the relative amount of variance explained, we devised a subsampling scheme where we randomly selected 15 bromeliads from a randomly selected field visit within a bioclimatic zone (18 zones, Figure 1). Note that the minimum number of bromeliads per site is 18 (Table A1), therefore for some sites, most bromeliads are selected in every sub-sampling procedure. We found that 15 bromeliads is the minimum number that still provides a comprehensive sample of the community within a field visit. Every time we performed this sub-sampling procedure, we ran the multivariate statistical analysis and compiled the main results (sum of squares). We repeated this process 1000 times. We used the marginal sum of squares in the analysis without interactions and the sequential sum of squares in the analysis with interactions to the variation explained by the main effects. From these runs, we obtained a distribution of P-values and sums of squares. We do not report P-vallues of individual runs, because they do not represent valid independents tests, however, we do report the P-values of a nonsubsampled analysis in the appendix (Tables A3 and A4). Some of the distributions of sums of squares were skewed, while some were normally distributed (Figure A5). To summarize this variation, we first calculated the mean of the sum of squares explained by each predictor across all subsamplings, and then calculated the total contribution of local conditions, climate variables, and their interaction. This procedure allowed us to take advantage of the central limit theorem to ensure that the addition occurs on normally distributed means.

**Figure 1:**
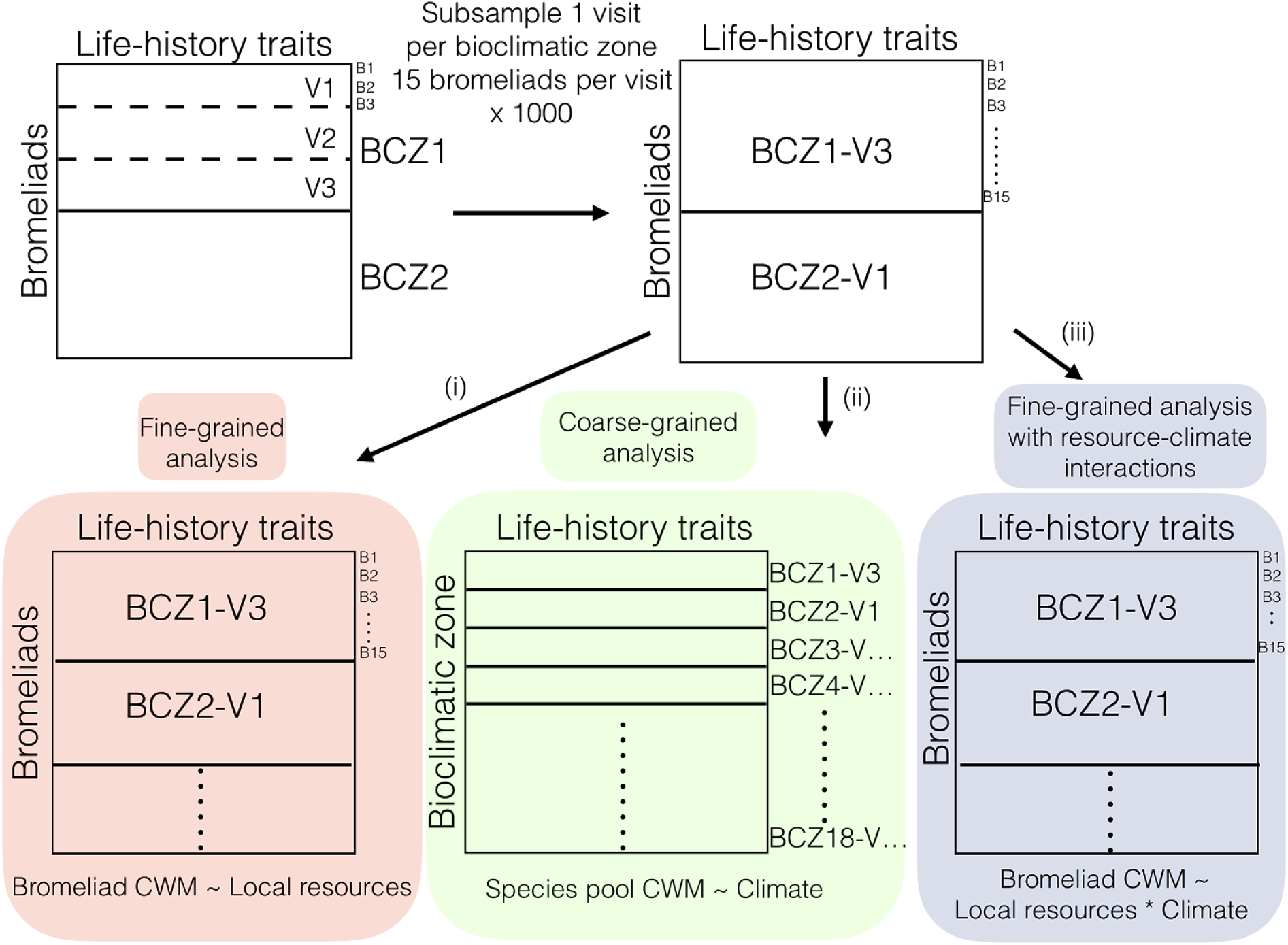
Schematic representation of the analysis. We tested for the effect of environmental conditions on trait composition in three main analyses. BCZ1, BCZ2, BCZ… represent the bioclimatic zone, which can contain multiple visits V1, V2, V3, V… We sampled one visit for each bioclimatic zone represented by BCZ1-V3, that is, bioclimatic zone 1, visit 3. (i) At the fine grained (red area) analysis, we tested for the effect of local conditions on bromeliad weighted means (CWM). (ii) At the coarse-grained analysis (green area), we tested for the effect of climate on species pools CWM. (iii) We tested the interaction between climate and local conditions in determining bromeliad CWM (blue-gray area).

We performed the multivariate analysis of variance for each spatial scale to replicate a ‘fine-grained analysis’ (Figure 1a), a ‘coarse-grained analysis’ (Figure 1b) and a fine-grained analysis with resource-climate interactions (Figure 1c). Thus, we are able to compare the explanatory power of these models to explain functional trait composition if we only had local conditions, climate information or both, (i) For the fine-grained analysis, we used each bromeliad as the sampling unit and only tested the effects of local conditions (that is the environmental conditions that were measured for every bromeliad: water level, detritus amount, and canopy cover). We restricted the sub-sampling to field visits (site by year combination) to ensure that maooinvertebrate traits relating to species that only occurred in a single bioclimatic zone (most species) were not mixed between countries or years (Figure 1 - finegrained analysis). Analysis using bromeliads as sampling units and only climatic variables as predictors were used to filter the 19 bioclimatic variables to a smaller subset. We retained climatic variables that explained a significant proportion of variation in at least 5% of the runs (BC2, 4,15 and 17), and which were then used in subsequent analyses. In 1(000 randomizations we expect at least 5% of runs to appear significant by chance (type I error), so we only report explanatory variables that are significant in >5% of runs, (ii) For the coarse-grained analysis, we used the ‘bioclimatic zone’ as units for which we calculated the species pool CWM by summing the abundance of all the morphospecies across the subsampled 15 bromeliads and only tested the effects of climatic variables (Figure 1 - coarse-grained analysis). (H) To test for the interactions between climate and local conditions, we used the bromeliad as the sampling unit, and tested the effect of local conditions, climate and their interactions (Figure 1 - finegrained analysis with resourceclimate interactions). We did not include the interaction between canopy cover and climatic conditions because few bioclimatic zones had both open and closed canopy, and consequently canopy cover would be confounded with bioclimatic zone.

All multivariate analyses were performed using the vegan package (Oksanen et al. 2017). Mixed effect models were performed using *Ime4* R package (Bates et al. 2015) and all analyses were done using the R programming language (R Core Team 2016). The code for the sub-sampling and statistical analysis, as well as the adaptation of the PERMANOVA can be found in: https://github.com/lmquzman/Climate_invertebrate_traits.

## Results

### Spatial scale of environmental variation

We determined the spatial scale of variation in our three local conditions: total detritus, water volume at time of sampling, and whether the canopy was open or closed. This analysis gave some indication of the potential power of each variable to explain variation in synthetic trait composition (i.e. little variation at a given scale indicates a lower likelihood of a significant effect at that given scale). Variation in total detritus was greatest at the level of the bioclimatic zone (39.2% of variation) but also was high at the local scale of the field site (31%). Variation in water volume was greatest at the local scale of field visits (42.6%) and minimal at the level of bioclimatic zone (2.7%). Finally, variation in canopy cover was highest at the level of bioclimatic zone (73.5%) and lowest at the level of field visit (26%) (Figure A3). We also found that these three local conditions were only weakly correlated across and within zones. The correlation values across bioclimatic zones ranged between −0.01 and 0.35, while the correlation values within bioclimatic zones ranged between −0.32 and 0.28 (Table A2).

### Fine-grained analysis

Invertebrate traits varied among every bromeliad within a field visit (Figure 2). We found that only a small amount of trait variation in CWMs (6.1%) could be explained by local conditions, and that no single local condition explained most of this variation. The amount of variation explained ranged from zero to 19.7% depending on the subset of bromeliads selected, and the distribution of variation explained was skewed (Figure A1, Table 1 - Finegrained analysis).

**Figure 2:**
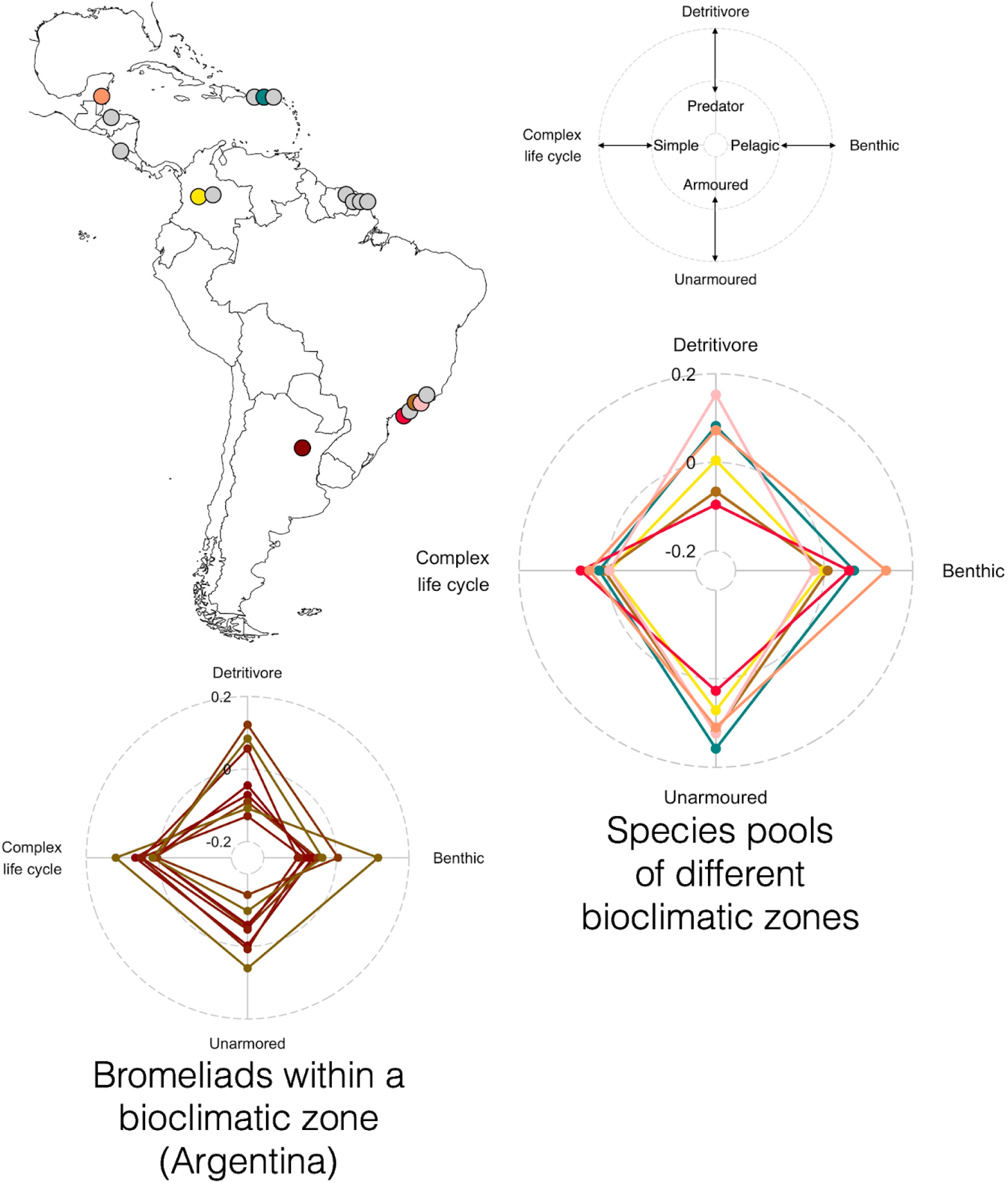
Invertebrate traits at the level of bromeliads and at the level of species pools, for bromeliads sampled in bioclimatic zones throughout the Neotropics (map, top left). Empty spider plot (top right) shows all the four trait axes and their directions, and forms a key for the two filled spider plots (bottom). Filled spider plots summarize the CWM of the four trait axes in a single bromeliad (bottom left) and the CWM in some example species pools (bottom right). Colours on spider plots correspond to bioclimatic zones on map, with those zones not shown in spider plots indicated in grey on the map.

**Figure 3:**
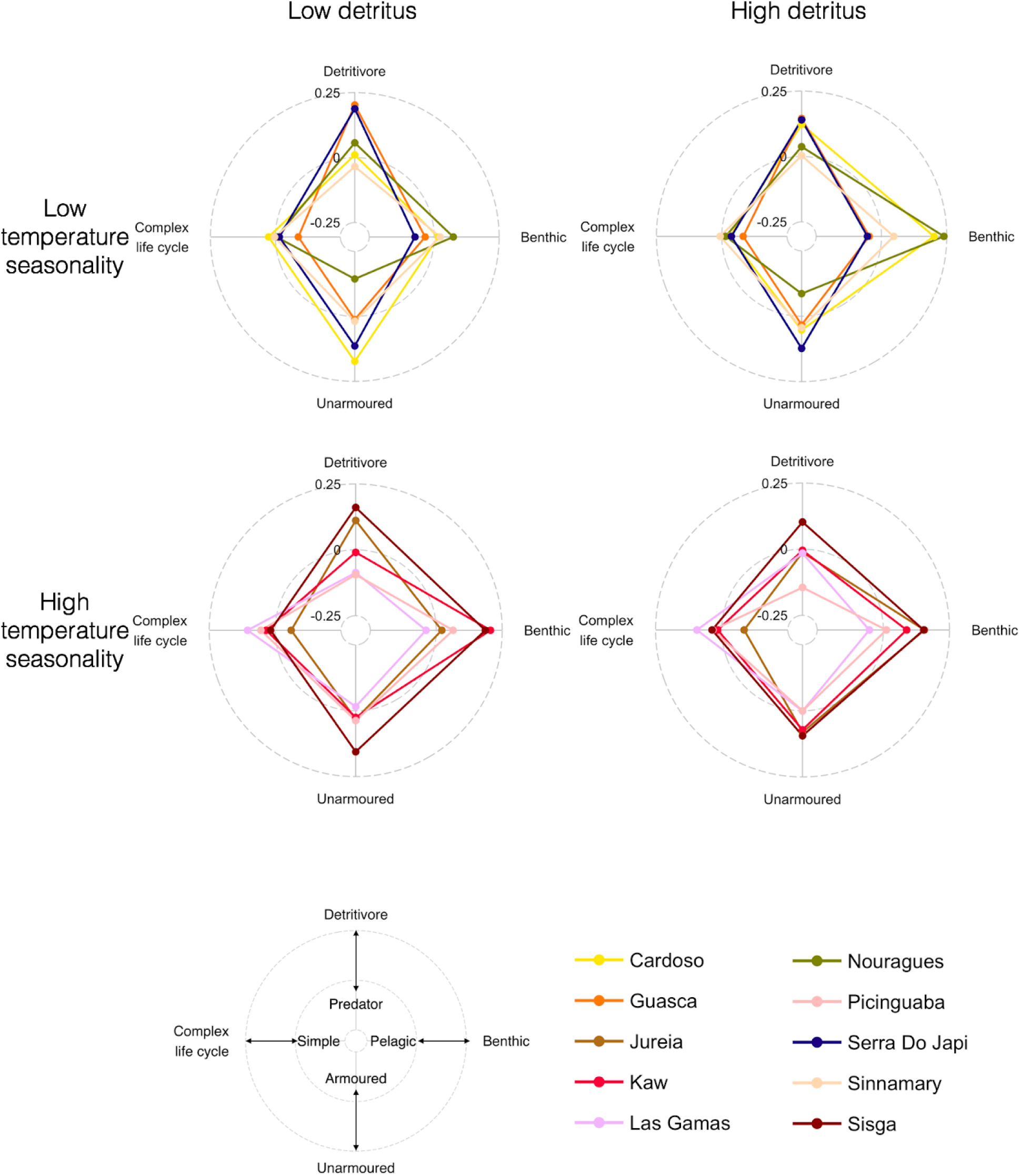
Bromeliads are characterized by their community-weighted mean traits for bioclimatic zones with low and high detritus, and high and low temperature seasonality. Detritus affects particularly the Benthic-Pelagic axis: bromeliads with high detritus have more benthic organisms (i.e. high Benthic-Pelagic axis values). Bioclimatic zones with high temperature seasonality have more organisms with complex life cycles (i.e. higher values in the Complex life cycle - Simple axis).

**Table 1.** Synthetic trait composition (CWM) explained by local conditions in the finegrained analysis and by climatic variables in the coarsegrained analysis. The analysis using the local conditions uses the CWM for each bromeliad. This analysis is blocked within each bioclimatic zone. The analysis using the biogeographic climatic variables uses the CWM for the species pool for each bioclimatic zone.

| Fine-grained analysis |  |  |
| --- | --- | --- |
| Predictor | Percentage of total sum of squares |  |
|  | Mean | Median |
| Canopy cover | 2.68 | 2.19 |
| Detritus | 2.30 | 1.92 |
| Water volume | 1.18 | 0.99 |
| Full model | 6.16 | 5.1 |
| Coarse-grained analysis |  |  |
| Mean Diurnal Temperature Range | 14.7 | 14.7 |
| Temperature Seasonality | 10.2 | 10.1 |
| Precipitation of Driest Quarter | 7.64 | 7.69 |
| Precipitation Seasonality | 7.39 | 7.25 |
| Full model | 39.93 | 39.44 |

### Coarse-grained analysis

Species pools differed in the relative proportions of invertebrate traits (Figure 2). In the coarse-grained analysis we found that climatic variables explained on average 39.9% of the variation in the trait composition of species pools (Table 1 - Coarse-grained analysis). The range of explained variation was large (14 to 47%) depending on the subset of bromeliads selected (Figure A2). The amount of variation explained in this analysis is not necessarily directly comparable to that in the fine-grained analysis because the scale of the response variable (CWMs) is different; for this analysis, we aggregated species at the site rather than bromeliad level. This aggregation changes the mean by weighting bromeliads with more individuals more heavily, and also reduces the number of observations, which tends to raise the R2 values. Four out of 19 bioclimatic variables explained substantial variation in trait composition of the macroinvertebrates across the Neotropics: mean diurnal range in temperature (BC2), temperature annual seasonality (BC4), precipitation annual seasonality (BC15), and precipitation of the driest quarter (BC17) (Figure A4; Table 1). Species pools from zones with high mean diurnal temperature range and high precipitation seasonality tended to be dominated by armoured invertebrates (Figure A5d, A7d). These climatic variables also differed in their effect on trophic traits: detritivores were favoured in zones with high precipitation in the driest quarter (Figure A8c), whereas predators were favoured in zones with high precipitation seasonality and mean diurnal range (Figures A5c, A7c).

### Fine-grained analysis with resource-climate interactions

The full model - using both the climatic and local resource environmental gradients to explain traits within individual bromeliads - explained between 27.2 and 441.1% of trait variation, when all the explanatory variables were included, with an average of 36.5% of the variation explained (Figure A9). We found that the local conditions explained 8.7%, climate explained 17.7%, and their interaction explained 10% of the variation in community weighted functional traits (CWMs) on average across all runs. Among the local conditions tested, detritus explained more variation than canopy cover or water volume. Of the climatic conditions tested, mean diurnal range in temperature (BC2) explained more variation than did the other climatic variables. Bromeliads with high mean diurnal range in temperature typical ly had more complex and unarmoured invertebrates (Figure A10). The crossscale interaction that explained the most variation was detritus amount and temperature seasonality (BC4; Figure 3–4, Figures A11, Table 2). Specifically, detritusrich bromeliads in zones with seasonal temperatures tended to contain more unarmoured invertebrates, predators, and invertebrates with complex life cycles (Figure 4). No single explanatory variable consistently expained the most variation in CWMs, rather, each variable contributed a small amount to the total amount of variation explained by the full model, which taken together explained more than a third of the variation in functional traits.

**Table 2:** Synthetic trait composition explained by local conditions, biogeographic climatic variables and their interactions. This analysis used the CWM for each bromeliad. We did not include the interaction between canopy cover and climatic conditions because few bioclimatic zones had both open and closed canopy, therefore canopy cover would be confounded with bioclimatic zone.

|  | Percentage of total sum of squares |  |
| --- | --- | --- |
|  | Mean | Median |
| *Local conditions* |  |  |
| Canopy cover | 2.43 | 1.86 |
| Detritus | 4.95 | 4.86 |
| Water volume | 1.35 | 1.09 |
| *Climatic variables* |  |  |
| Mean Diurnal Temperature Range | 5.55 | 5.55 |
| Temperature Seasonality | 3.55 | 3.31 |
| Precipitation of Driest Quarter | 4.66 | 4.57 |
| Precipitation Seasonality | 3.97 | 3.99 |
| *Interactions between local conditions and climatic variables* |  |  |
| Detritus : Mean Diurnal Temperature Range | 1.23 | 1.0 |
| Detritus : Temperature Seasonality | 2.22 | 2.12 |
| Detritus : Precipitation of Driest Quarter | 1.33 | 1.25 |
| Detritus : Precipitation Seasonality | 1.56 | 1.39 |
| Water volume: Mean Diurnal Temperature Range | 0.84 | 0.68 |
| Water volume: Temperature Seasonality | 0.95 | 0.86 |
| Water volume: Precipitation of Driest Quarter | 1.37 | 1.20 |
| Water volume: Precipitation Seasonality | 0.52 | 0.44 |
| *Total sum of squares* |  |  |
| Full model | 36.49 | 36.62 |

**Figure 4:**
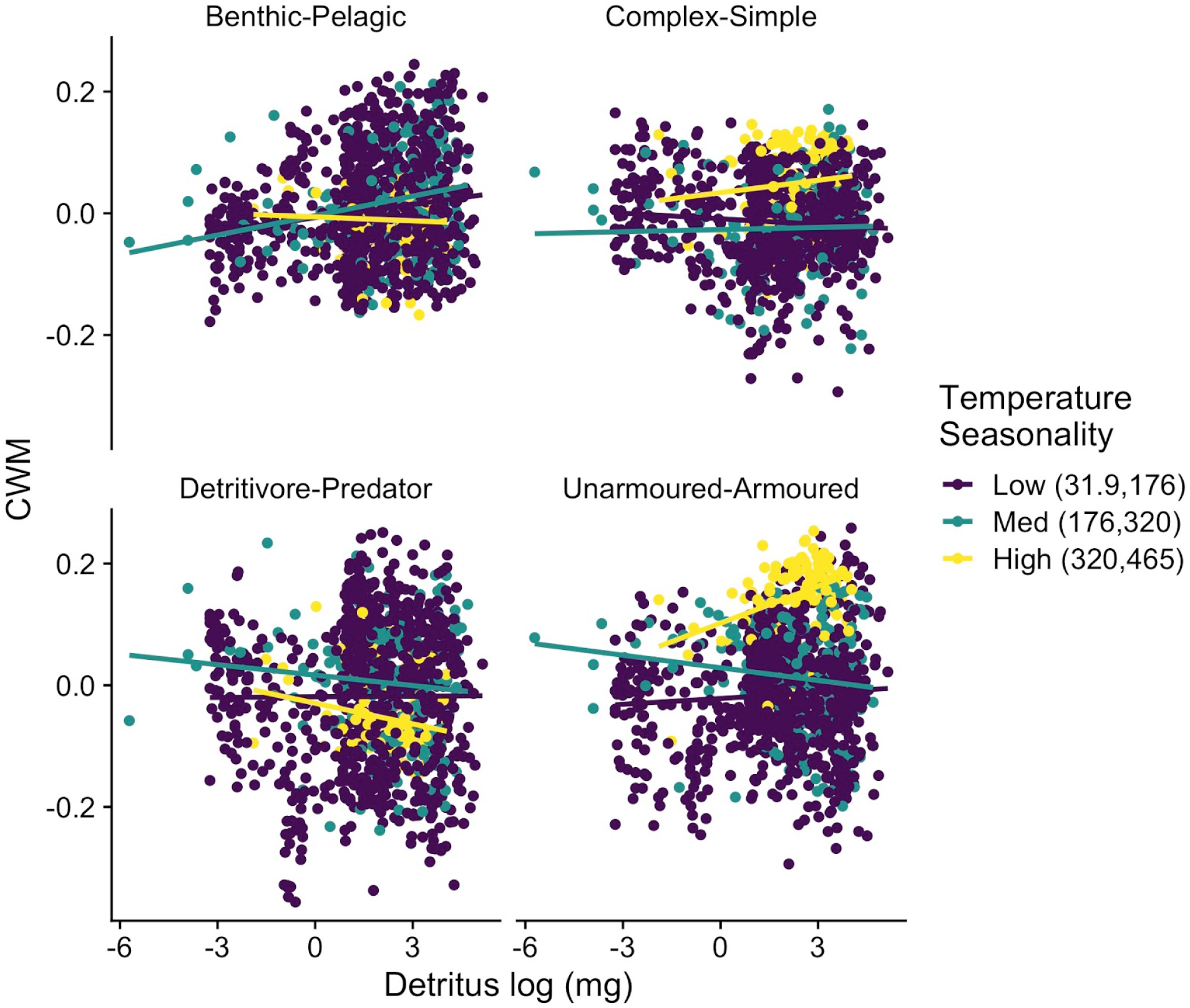
The four trait axes differ in their relationship with the total amount of detritus which is also mediated by temperature seasonality. Each point is the community weighted mean of a single bromeliad. The coloured lines are simple linear regressions intended only to improve the visualization of the data and not meant to be used for formal analysis since the CWM are multivariate.

## Discussion

Our analyses demonstrate the importance of climate and cross-scale interactions of climate with local conditions on the functional traits of macroinvertebrate communities. Climate explained 40% of the trait variation in regional species pools, corroborating the hypothesis that trait composition is primarily expained by the environment at the biogeographic scale. Climate also explained a substantial amount (28%) of trait variation at the scale of individual bromeliads, both directly (18%), and by modifying the response of traits to local conditions in cross-scale interactions (10%) (Figure A9). By contrast, local conditions of bromeliads, by themselves, explained only 6-9% of trait variation depending on the model used. The fact that we were able to explain trait variation with both climate and local conditions means that functional traits are not entirely determined by stochastic factors, historical contingency and phylogenetic constraints. Although stochastic and biogeographic factors no doubt affect trait distribution, they are not strong enough to completely overwhelm or erase the effects that climate and local conditions have on the functional traits of bromeliad macroinvertebrate communities. The implication is that local adaptation and/or filtering occurs in response to both local and climatic conditions, and ultimately shapes the ecological communities of bromeliad macroinvertebrates and their suite of traits.

The most important climatic factor in explaining trait variation was mean diurnal range of temperature. Combinations of precipitation and temperature were also important.

Detritus and canopy cover were the most important local conditions, but only explained 6.1 % of the variation in invertebrate trait composition. In general, not a single predictor (either local conditions or climate) explained very large amounts of variation in trait composition. Instead, the aggregated effects of each predictor contributed to the total variation explained.

Our fine-grained analysis with cross-scale interactions allowed us to conclude that climate explains more variation in macroinvertebrate trait composition than local conditions. There are several possible explanations for this pattern. First, our synthetic trait axes may have captured fundamental differences in the strategies of species for tolerating climate-related stress, but not for exploiting local habitat heterogeneity. One of the most important stresses in the bromeliad system is hydrologic variability, including periods of drought. Some species are able to withstand drought with drought-resistant eggs (e.g. Wyeomyia spp.: Dézerald et al. 2015), whereas others have drought-tolerant larvae (e.g. Tipulidae larvae: Amundrud and Srivastava 2015). Many mosquito larvae are sensitive to drought because their legless larvae require water to move (e.g. Culex spp.: Amundrud and Srivastava 2015). However, odonates - a dominant predator in the food web - are vulnerable to drought because of their long larval stage (Guzman et al. 2019, Srivastava et al. 2020a). Therefore, multiple trait axes, as used in our study, are needed to capture traits relevant to drought tolerance, including resistant life forms, larval duration (i.e. cohort production interval), and pelagic requirements. Geographic patterns in drought predict the distribution of invertebrate families that comprise the species pool of bromeliad invertebrates (Srivastava et al. 2020b) and families often have unique functional traits (Céréghino et al. 2018). Climate is likely a better predictor of such geographic patterns in drought prevalence than bromeliad water volume, as water volume is only measured on the particular day of sampling and is very dynamic between days. Invertebrate mortality rarely follows a single day of drought; instead mortality ranges from 111-73% after 18 days without water (Amundrud and Srivastava 2015). Similarly, experimental exclusion of rainfall from French Guiana bromeliads led to changes in functional trait composition of invertebrates only after six weeks without rain (Dézerald et al. 2015). Although no single climate variable dominated the effects on traits, many of the top climate variables were related to variation (daily or seasonal) in temperature and precipitation as might be anticipated if climate affected traits via drought prevalence. Our conclusion that temperature seasonality was an important determinant of trait composition is similar to Swenson et al. (2012).

A second explanation is that deterministic filtering by local conditions is largely overwhelmed by stochasticity in the colonization and emergence rates of invertebrates from bromeliads. The majority of invertebrates in bromeliads are insects and thus have complex life cycles, meaning that only the egg to larval or pupal stages are aquatic. Larval development can be as fast as two weeks for mosquitoes, and the majority of insects (except odonates) have cohort production intervals of less than 30 days (Dézerald et al. 2017). This is a relatively short period for the amount of detritus, water or light to limit their abundances, and suggests that abundances may be more affected by oviposition and predation - both of which have an important stochastic component. Furthermore, because low abundances of species in bromeliads can indicate either insufficient oviposition, successful completion of the larval stage and emergence, or larval mortality, even deterministic effects of local factors may result in a complex array of positive and negative effects on abundance. Given that the population dynamics of species with complex life cycles (i.e. insects) occurs at scales larger than the bromeliad (LeCraw et al. 2014), we might expect stronger trait-environment correlations to be found at these larger scales, scales which are based on the bioclimatic zone and /or the species pool.

A third possibility is that the suite of synthetic trait axes and local variables we used for analysis somehow predetermined greater matching of traits with climatic variables than with local conditions. However, both the traits and local conditions used in this study have been identified in previous studies as important factors determining community composition (Richardson 1999, Usseglio-Polatera et al. 2000, Dézerald et al. 2014). The four synthetic trait axes represent major fundamental niche dimensions such as trophic position, habitat, life history and defence (Céréghino et al. 2018), and explained 45% of the total variance in species traits from the 12 traits we assembled. Although the main goal of this study was to explain variations in those four main ecological strategies or four PCA axes, other important ecological strategies (PCA5, PCA6, PCA7, …)) could also be influenced by local and bioclimatic conditions, however, these other axes were not significant in (Céréghino et al. 2018)) and did not have biological interpretations. The four ecological strategies we studied here, have previously been identified as basic niche dimensions in other systems and in other clades (Winemiller et al. 2015), suggesting that they may be general across different types of communities, and perhaps, broadly applicable to aquatic invertebrates in other ecosystems. Extensive research on bromeliad communities has demonstrated that local conditions such as water volume, detrital amount and canopy cover affects bromeliad community structure, including predatonprey ratios and species richness in bromeliads (Richardson et al. 2000, Srivastava et al. 2008, Dézerald et al. 2014). There are well-known mechanistic reasons behind these relationships. The amount of light available to a bromeliad (Le. canopy cover) determines algal productivity, and therefore, the relative importance of detritus versus algae in the diet of different macroinvertebrates (Farjalla et al. 2016). In general, detritus is the main source of nutrients in the bromeliad food web, and its quantity is related to overall macroinvertebrate biomass (Richardson et al. 2000). The amount of water found in a bromeliad at the time of sampling is related to the amount of habitat available to invertebrates, the risk of drought, and whether it is colonized by predators, and as such, habitat size is an important predictor of species richness, species composition and trophic structure (Srivastava et al. 2008, Amundrud and Srivastava 2015, Petermann et al. 2015). In an experiment where many of these factors were controlled for, local variation in rainfall impacted the community structure of bromeliad macroinvertebrates (Srivastava 2020b)), and in extreme cases, caused an inversion of the trophic pyramid (Romero et al. 2020)). So a combination of local conditions will have some effect on community dynamics and the distribution of traits.

Our conclusion that climate overwhelms local conditions in driving community trait structure contrasts to studies on plant communities by Bruelheide et al. (2018) who found that microenvironmental gradients were more influential than climate. This may be because two of the three local scale factors we analyzed varied more at biogeographical than local scales (Table A2).

An important conclusion of our analysis is that there are cross-scale interactions between environmental drivers of trait composition. That is, the effect of local conditions depends on the regional climate. Specifically, the effect of either detrital amount or water volume depended on temporal variation in precipitation and temperature at the field site. This may reflect the ability of large detrital-filled bromeliads to buffer the effects of climate variation on drought prevalence (Sirivastava et al. 2020a). Studies of functional traits that use coarse-grained data such as range maps or remote sensing data cannot test for such crossscale effects of the response of the community to local and climatic conditions. However, there is a growing number of fine-grained datasets with a complete census of the community for which interactions between local conditions and larger scale environmental constraints can be tested. Such datasets are particularly well represented by plants (e.g. Blonder 2018, Bruelheide et al. 2018), but also freshwater invertebrates (e.g. Aspin et al. 2019), fish (e.g. Winemiller et al. 2015), intertidal organisms (e.g. Menge et al. 1999) and marine coastal fishes (e.g. Hemingson and Bellwood 2018). The challenge for analyzing cross-scale effects in these studies is not the large-scale climatic data, but rather the fine-grained environmental data that matters for resource acquisition, competition, predation and facilitation. Fiine-grained microenvironmental data, only some of which was available in our study, is likely to be critical in determining the relative importance of environmental filtering and biotic interactions as well as the degree of context dependence (Blonder 2018). A particular advantage of our study was that we were able to measure the entire aquatic macroinvertebrate community at a fine scale at multiple locations across a wide biogeographic range, which then were assembled into a large database, through the support of the French data synthesis centre, CESAB. The randomized sub-sampling procedure was used to control for uneven sampling effort between sites, and gave robust estimates of variance explained between sites. Although subsampling reduces statistical power, we gained confidence in our among site comparisons.

Overall, we found that climate explained more variation than local conditions, and that the two scales interactively determine the functional traits of bromeliad macroinvertebrate communities across their Neotropical range. Our ability to contrast the effects of climatic vs. local conditions hinged on the compilation of multiple fine-grained datasets. We argue that in addition to sampling communities at local scales, ecologists should aggregate studies that span large ranges in climate variation in order to fully understand trait filtering at local, regional and global scales.

## Acknowledgements

This research is a product of the FunctionalWebs group funded by the synthesis center CESAB of the French Foundation for Research on Biodiversity (FRB; www.fondationbiodiversite.fr). We acknowledge the support provided by The Natural Sciences and Engineering Research Council of Canada (CGS-D) to L.M.G. and (Discovery Grant) to D.S.S., by UBC Four Year Fellowships to L.M.G., by the Agence Nationale de la Recherche through an Investissement d’Avenir grant (Labex CEBA, ANR-K0-LABX-25-011) to C.L. and R.C., by a BPE-FAPESP grant #201601209-9 to GQR, by CNPq-Brazil research grants to VDP (#307689/2014-0) and VFF (#312770/2014-6), by CONICET Piroyecto P-UE 22920160100043 and UNR AGR-290 to IMB, and by grants from the Royal Society of Edinburgh, the Carnegie Trust for the Universities of Scotland, acknowledge postdoctoral fellowship support from a PNPD-CAPES grant #20147/04603-4 to PJMdeO., a PNPD/CAPES grant #20130877 to N.A.C.M., a CNPq grant (#401345/2014-9) under the Ciências sem Fronteiras program to V.J.D and a FAPESP grant #20167/09699-5 to AZG. This is a publication of the Bromeliad Working Group.

