## Supplemental figures and tables for "Climate influences the response of community functional traits to local conditions"

**Appendix 1**

**Table A1:** Regions used with environmental conditions. BC2, Mean Diurnal Temperature Range; BC4, Temperature Annual Seasonality; BC15, Precipitation Annual Seasonality; BC17, Precipitation of the Driest Quarter.

| **Country** | **No. plants** | **Years sampled** | **Site** | **BC2** | **BC4** | **BC15** | **BC17** |
|---|---|---|---|---|---|---|---|
| Colombia | 36 | 2001 | Guasca | 8.50 | 32.29 | 58.76 | 130 |
| Colombia | 37 | 2000 | Sisga | 8.80 | 39.42 | 38.78 | 174 |
| Brazil | 18 | 2011 | Serra Do Japi Semidecidual | 9.72 | 227.68 | 64.37 | 120 |
| Puerto Rico | 70 | 1993, 1994, 1996, 2010 | El Verde Dwarf forest | 6.82 | 130.47 | 26.82 | 336 |
| Costa Rica | 106 | 1997, 2000, 2002, 2004, 2010 | Pitilla | 7.85 | 72.39 | 43.99 | 251 |
| Honduras | 157 | 2006, 2007 | Cusuco | 8.95 | 150.32 | 52.55 | 143 |
| Puerto Rico | 70 | 1993, 1994, 1996, 1997, 2010 | El Verde Palo Colorado | 8.16 | 132.60 | 25.98 | 335 |
| Puerto Rico | 90 | 1993, 1994, 1996, 1997, 2010, 2004 | El Verde Tabonuco Sonadora 400m | 8.87 | 144.59 | 27.40 | 328 |

| | | | | | | | |
|---|---|---|---|---|---|---|---|
| Brazil | 66 | 2008, 2011 | Cardoso | 7.32 | 287.02 | 45.29 | 277 |
| Brazil | 20 | 2013 | Jureia | 8.44 | 263.91 | 43.78 | 248 |
| French Guiana | 47 | 2008 | Kaw | 8.49 | 43.80 | 55.76 | 264 |
| Brazil | 70 | 2015 | Macae North | 8.49 | 203.25 | 47.81 | 114 |
| French Guiana | 199 | 2007, 2008, 2014 | Petit Saut | 8.08 | 48.09 | 44.86 | 317 |
| Brazil | 20 | 2011 | Picinguaba | 9.73 | 234.40 | 39.92 | 335 |
| French Guiana | 39 | 2011 | Sinnamary | 7.67 | 47.58 | 45.02 | 291 |
| French Guiana | 171 | 2006, 2009 | Nouragues | 9.41 | 47.79 | 50.34 | 309 |
| Argentina | 138 | 2004, 2005, 2010, 2012, 2013 | Las Gamas | 11.65 | 464.23 | 49.29 | 91 |
| Mexico | 40 | 2011 | Quintana Roo | 12.30 | 171.87 | 61.37 | 83 |

**Table A2:** Correlation of local conditions across and within bioclimatic zones.

| **Across bioclimatic zones** | | | |
|---|---|---|---|
| | Actual water volume | Total detritus | Canopy cover |

| | | | |
|---|---|---|---|
| Actual water volume | 1 | 0.33 | 0.35 |
| Total detritus | 0.33 | 1 | -0.013 |
| Canopy cover | 0.35 | -0.013 | 1 |
| **Mean correlation within bioclimatic zones** | | | |
| | Actual water volume | Total detritus | Canopy cover |
| Actual water volume | 1 | 0.29 | 0.19 |
| Total detritus | 0.29 | 1 | -0.32 |
| Canopy cover | 0.19 | -0.32 | 1 |

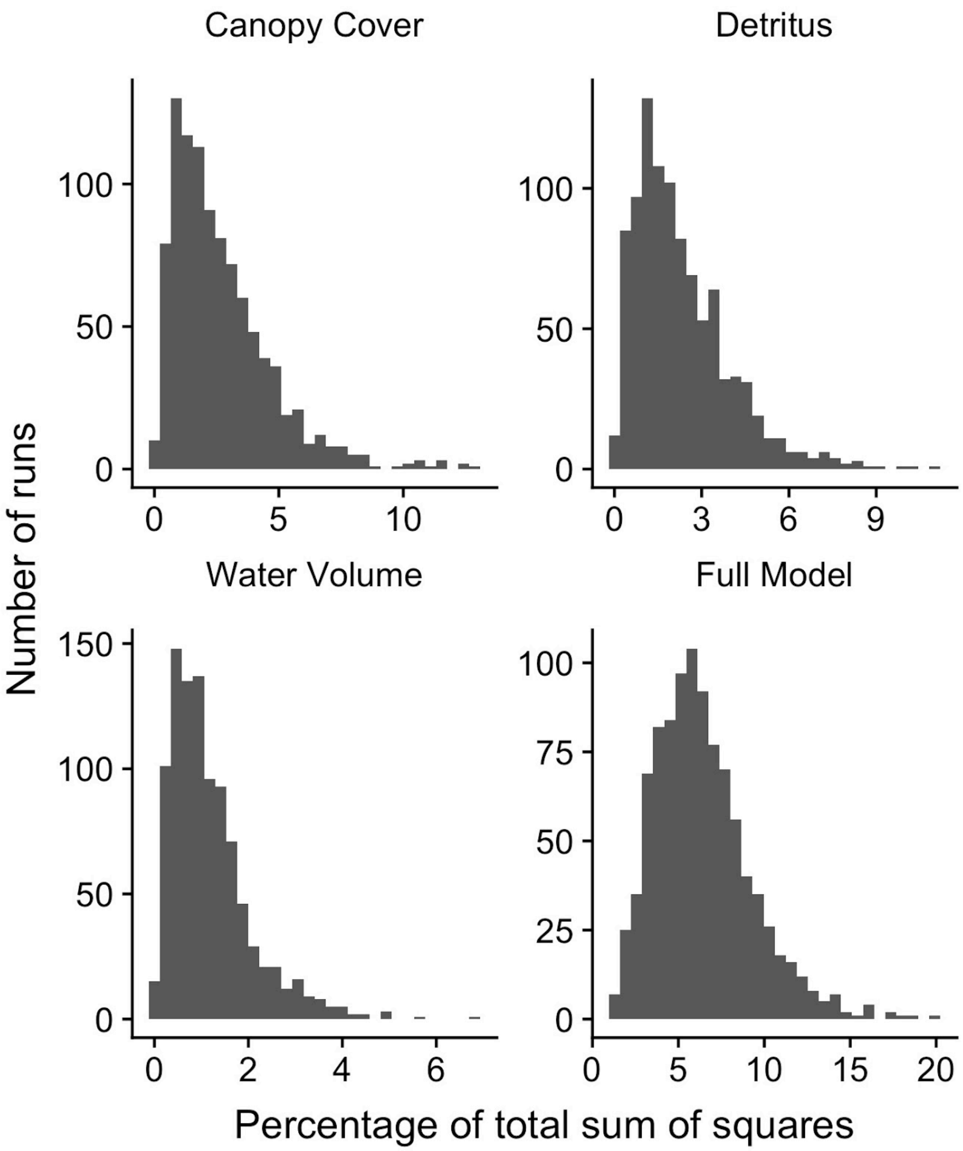


**Figure A1:** Distribution of the percentage of variation explained at the local scale (analysis i). In this analysis, we used each bromeliad as the sampling unit and only tested the effects of local conditions (that is the environment that can be measured for every bromeliad).

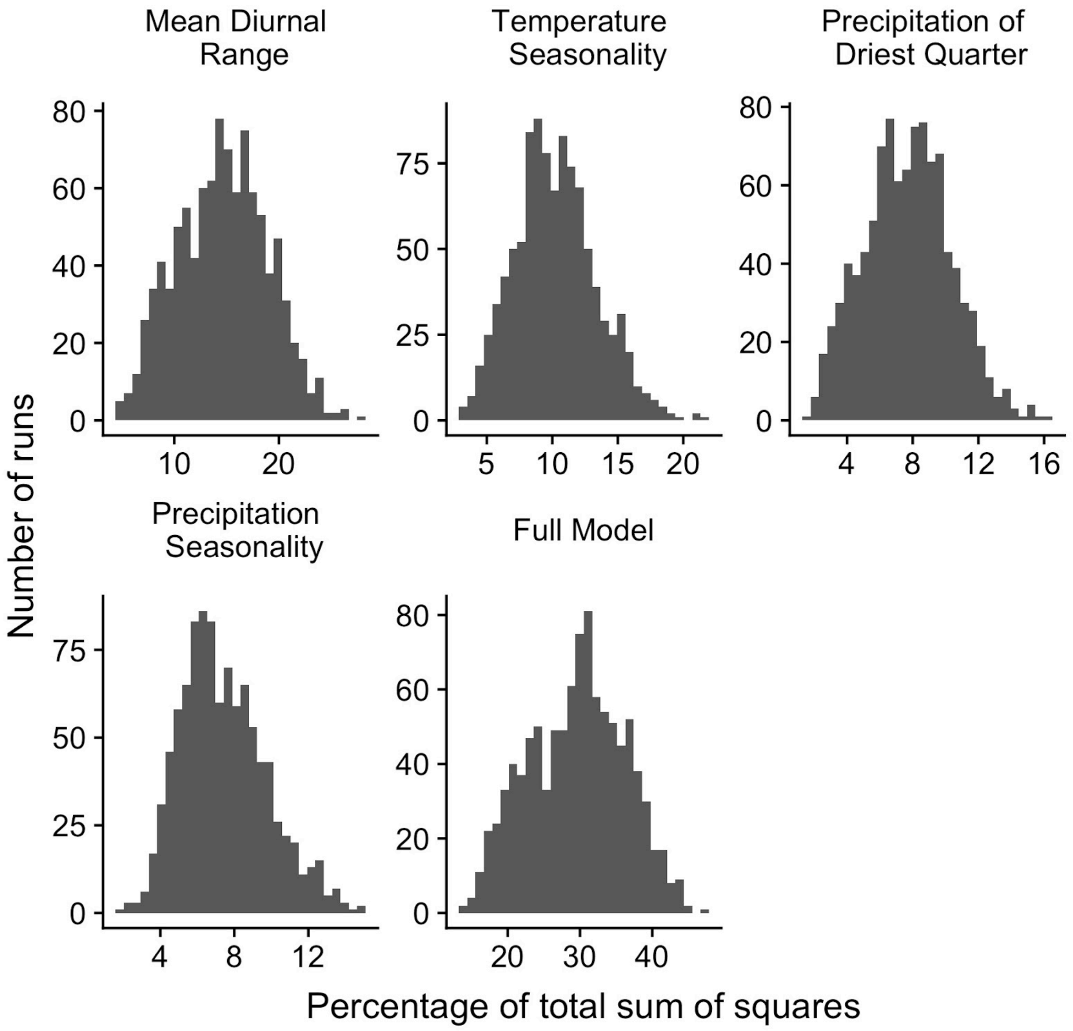


**Figure A2:** Distribution of the percentage of variation explained at the biogeographic scale (analysis ii). In this analysis, we used the ‘bioclimatic zone’ as the sampling unit and we calculated the species pool CWM by summing the abundance of all the morphospecies across the sub-sampled 15 bromeliads and only tested the effects of climatic variables.

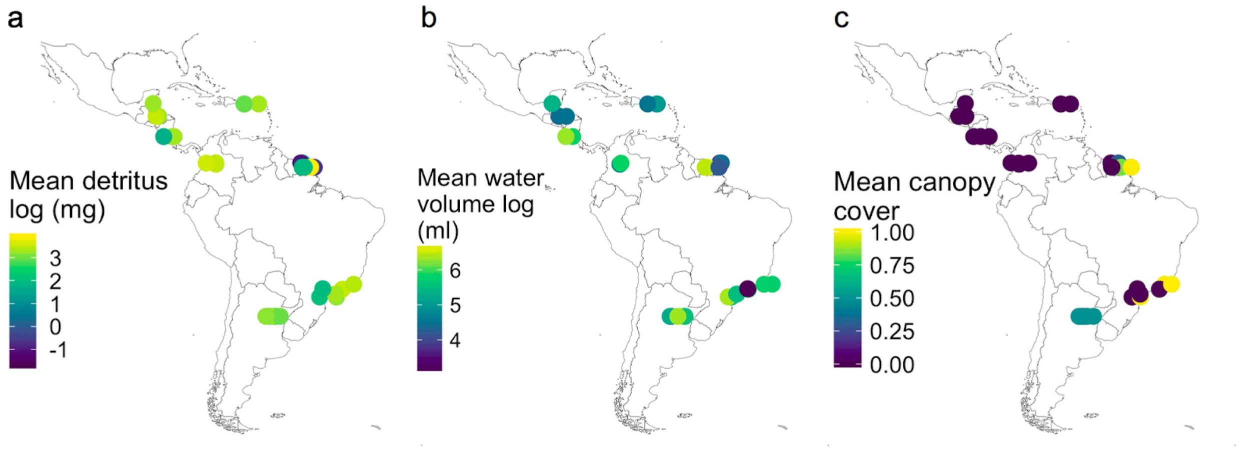


**Figure A3:** Variation in mean local conditions at 18 field bioclimatic zones: a) total detritus, b) maximum water volume, and c) canopy cover.

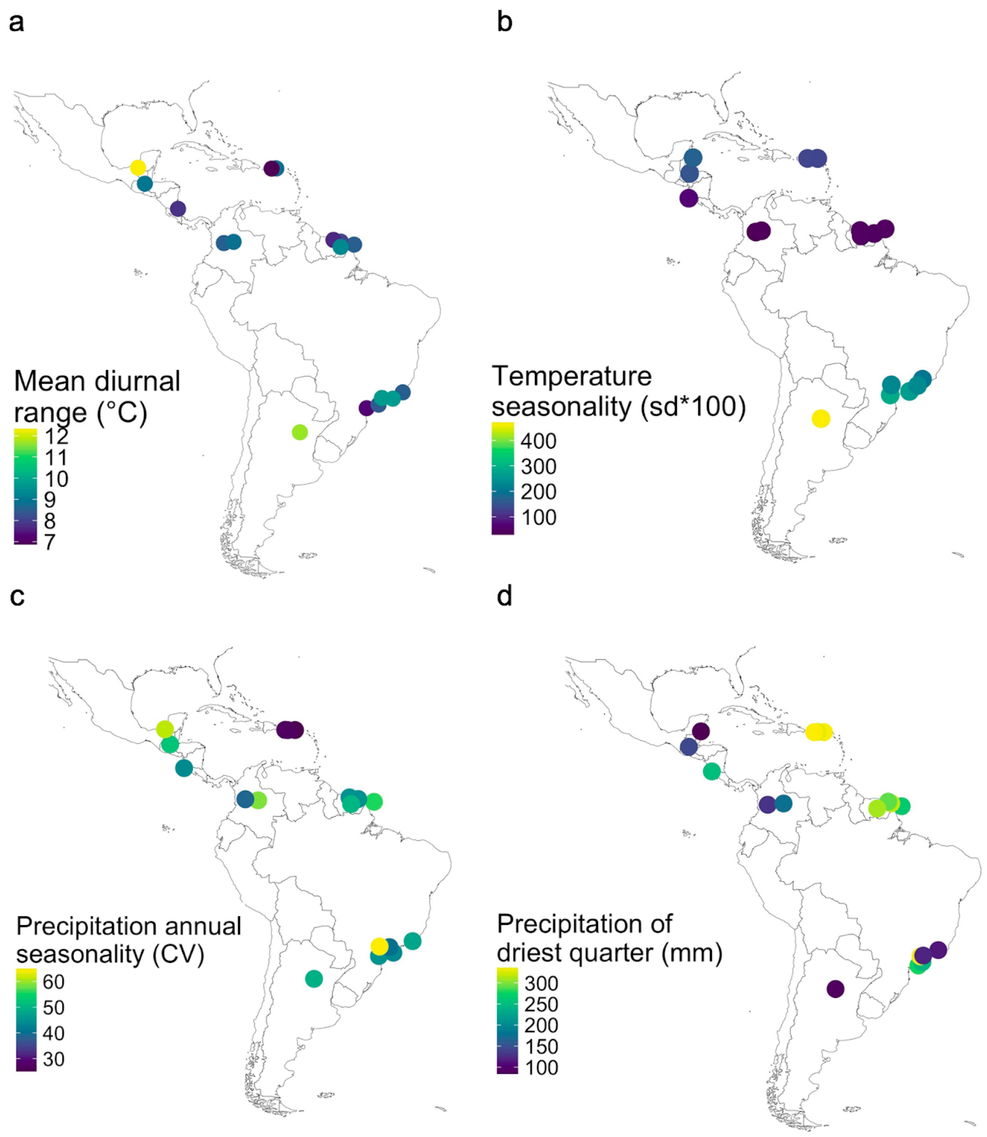


**Figure A4:** Bioclimatic conditions in each of the bioclimatic zones (colored dots). Only bioclimatic variables that were significant in more than 5% of the model runs are displayed: a) mean diurnal temperature range (BC2), b) temperature annual seasonality (BC4), c) precipitation annual seasonality (BC15), and d) precipitation of driest quarter (BC17).

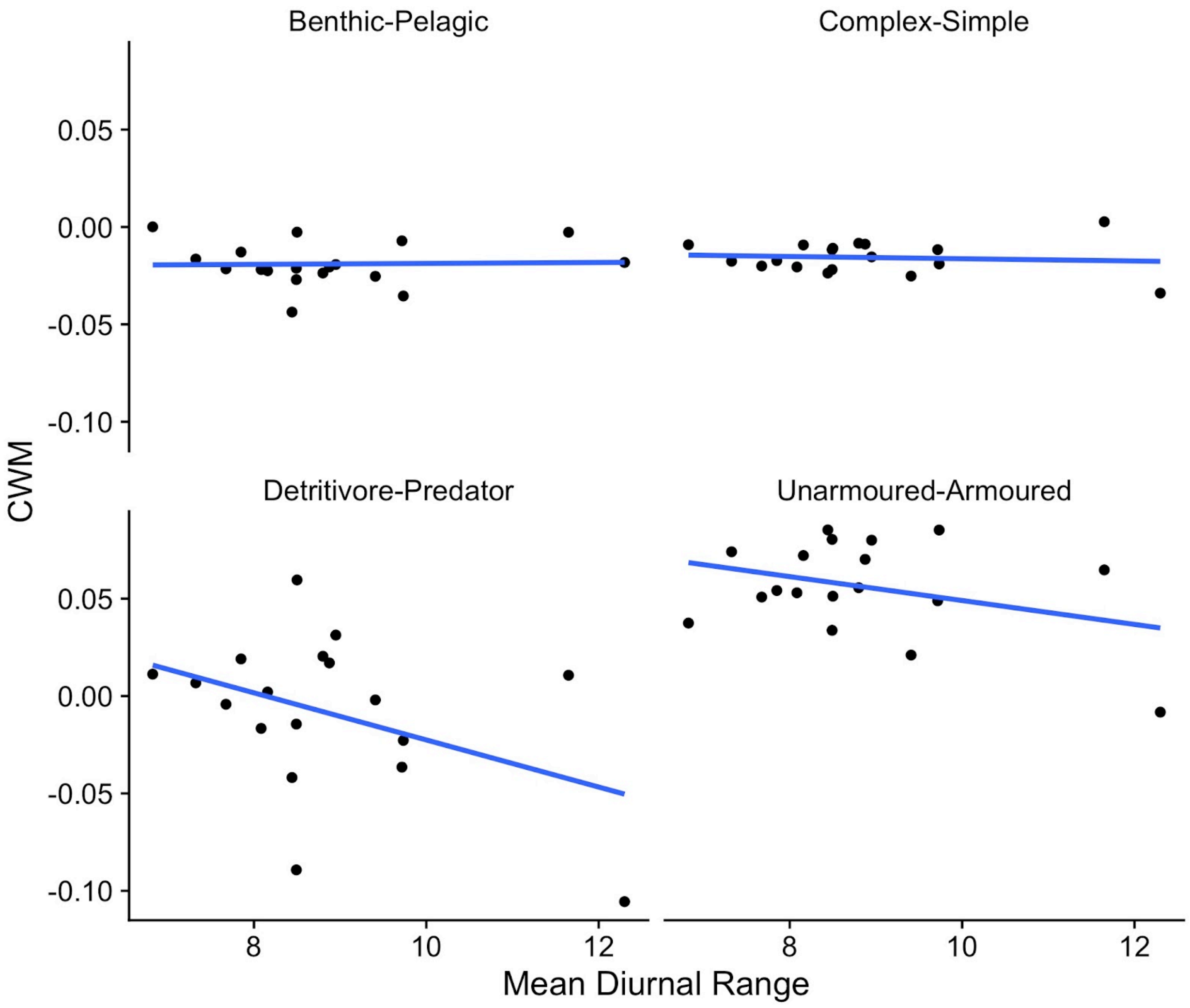


**Figure A5:** The four trait axes differ in their relationship with mean diurnal temperature range in a site. Each point is the mean community weighted mean of a bioclimatic region. Mean CWM was calculated by using the subsampling regime described in the methods and calculating the mean CWM across 1000 runs for each bioclimatic region. The blue lines are simple linear regressions intended only to improve the visualization of the data and not meant to be used for formal analysis since the CWM are multivariate.

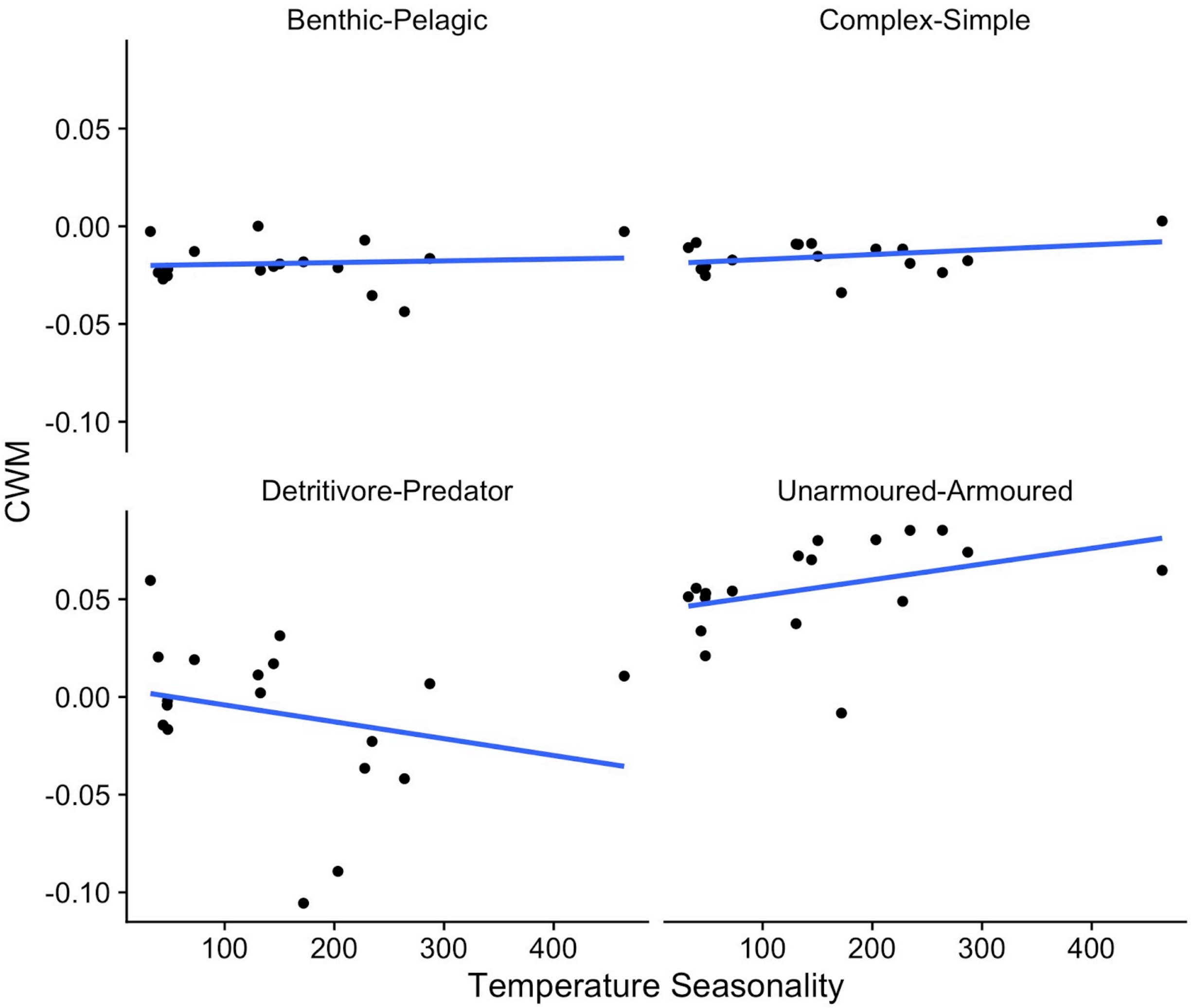


**Figure A6:** The four trait axes differ in their relationship with temperature seasonality. Each point is the mean community weighted mean of a bioclimatic region. Mean CWM was calculated by using the subsampling regime described in the methods and calculating the mean CWM across 1000 runs for each bioclimatic region. The blue lines are simple linear regressions intended only to improve the visualization of the data and not meant to be used for formal analysis since the CWM are multivariate.

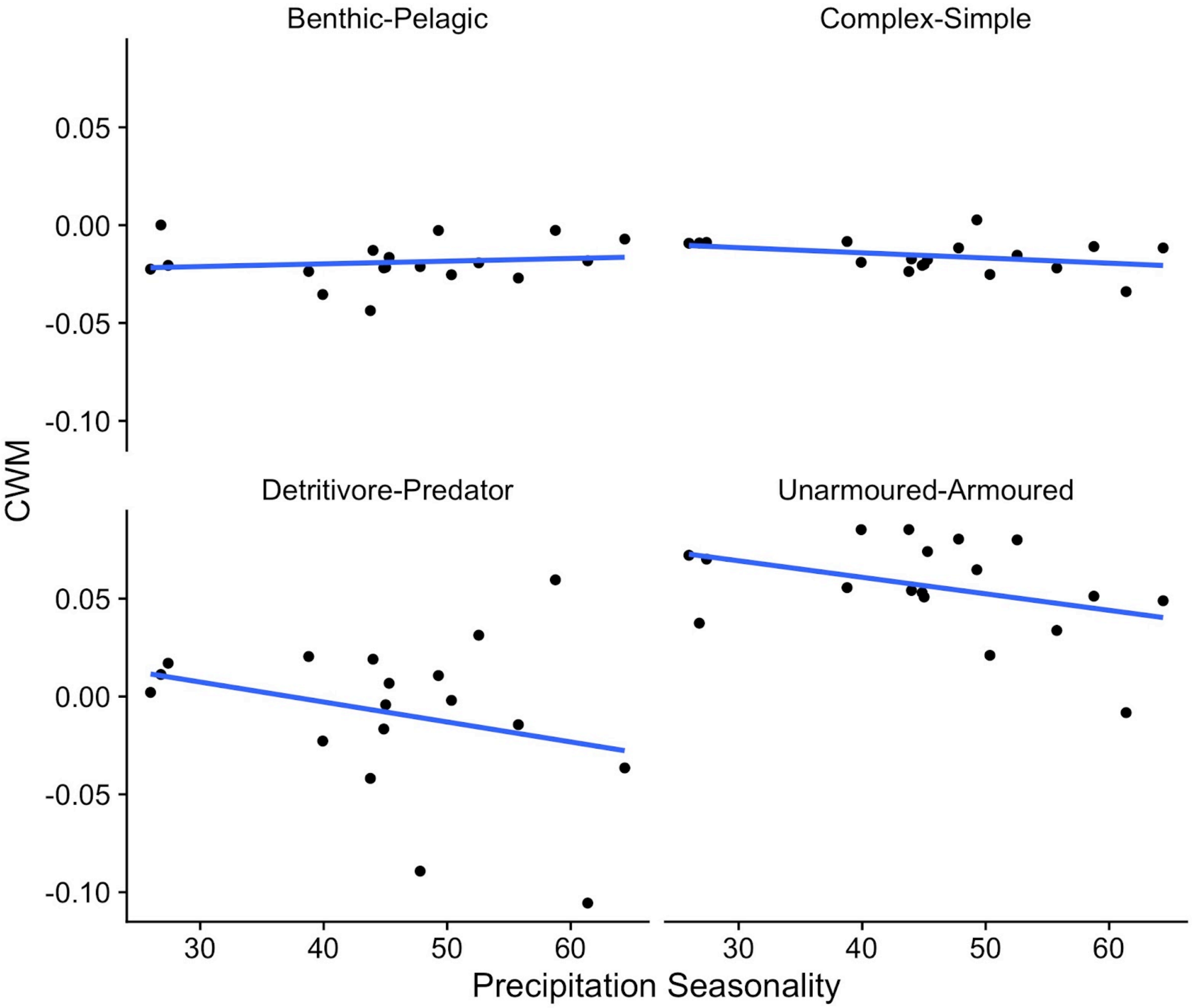


**Figure A7:** The four trait axes differ in their relationship with precipitation seasonality. Each point is the mean community weighted mean of a bioclimatic region. Mean CWM was calculated by using the subsampling regime described in the methods and calculating the mean CWM across 1000 runs for each bioclimatic region. The blue lines are simple linear regressions intended only to improve the visualization of the data and not meant to be used for formal analysis since the CWM are multivariate.

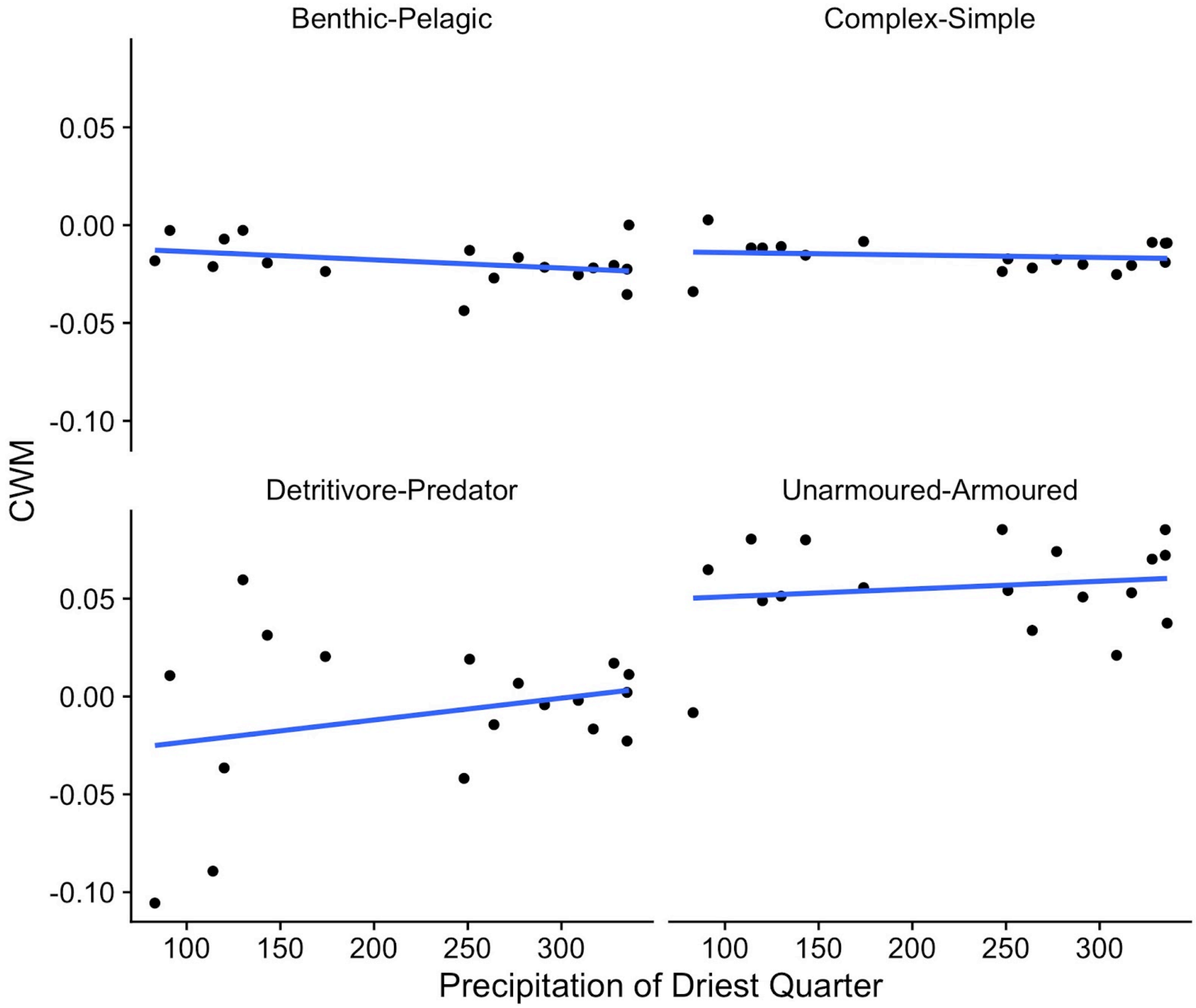


**Figure A8:** The four trait axes differ in their relationship with precipitation of the driest quarter. Each point is the mean community weighted mean of a bioclimatic region. Mean CWM was calculated by using the subsampling regime described in the methods and calculating the mean CWM across 1000 runs for each bioclimatic region. The blue lines are simple linear regressions intended only to improve the visualization of the data and not meant to be used for formal analysis since the CWM are multivariate.

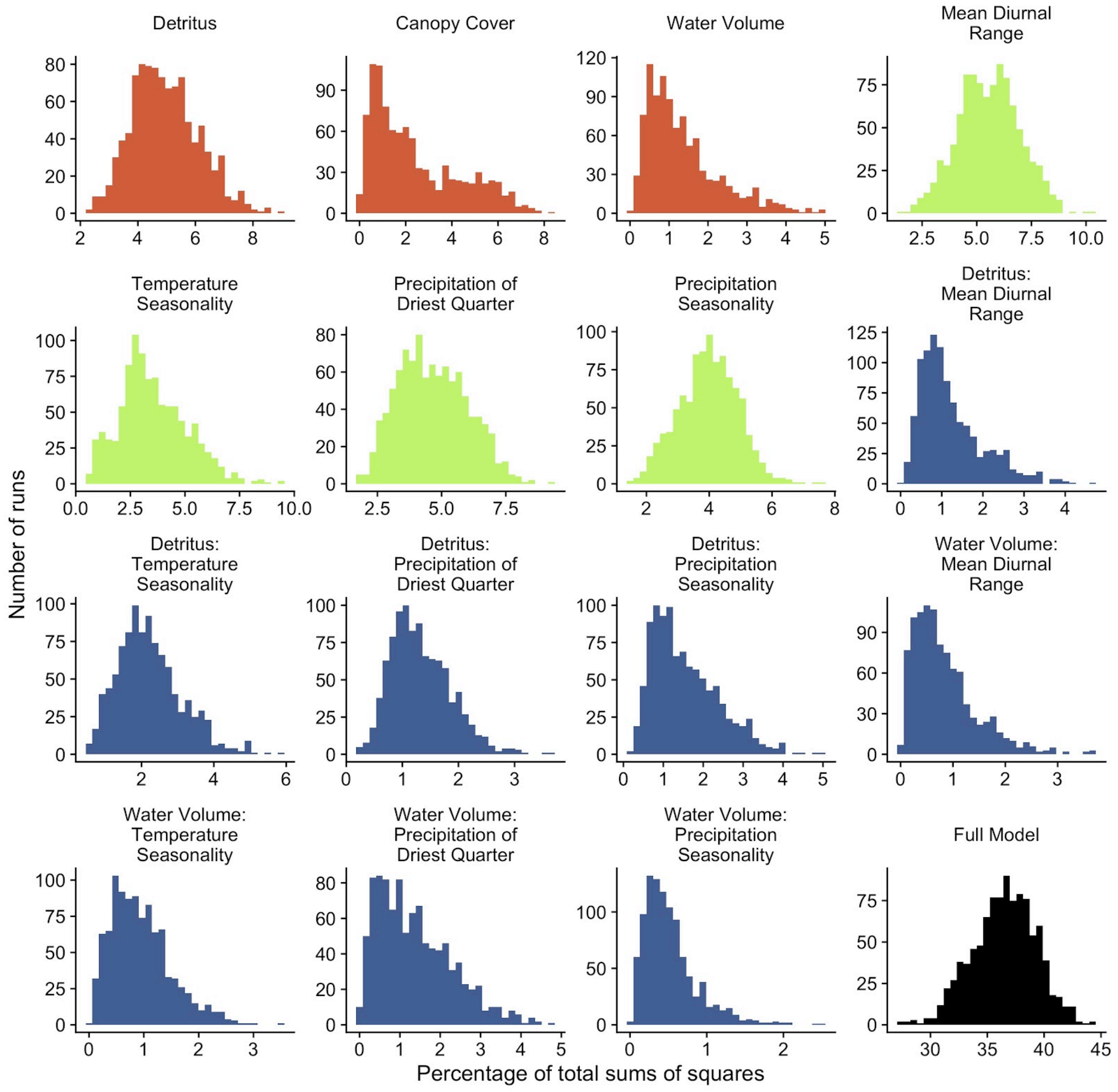


**Figure A9:** Distribution of the percentage of variation explained when we test for the effect of local conditions, local environmental conditions (orange), climate variables (green) and their interaction (blue) (analysis iii).

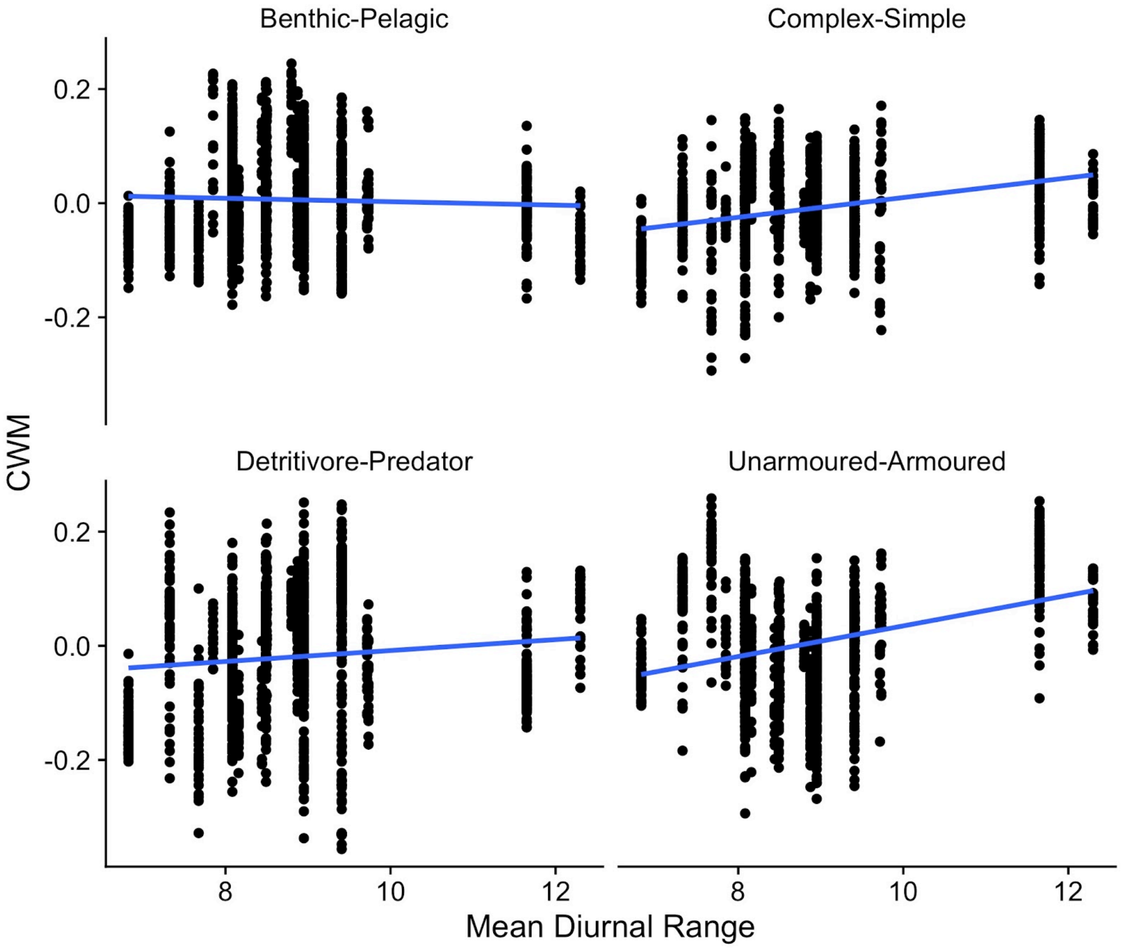


**Figure A10:** The four trait axes differ in their relationship with mean diurnal range. Each point is the community weighted mean of a single bromeliad. The blue lines are simple linear regressions intended only to improve the visualization of the data and not meant to be used for formal analysis since the CWM are multivariate.

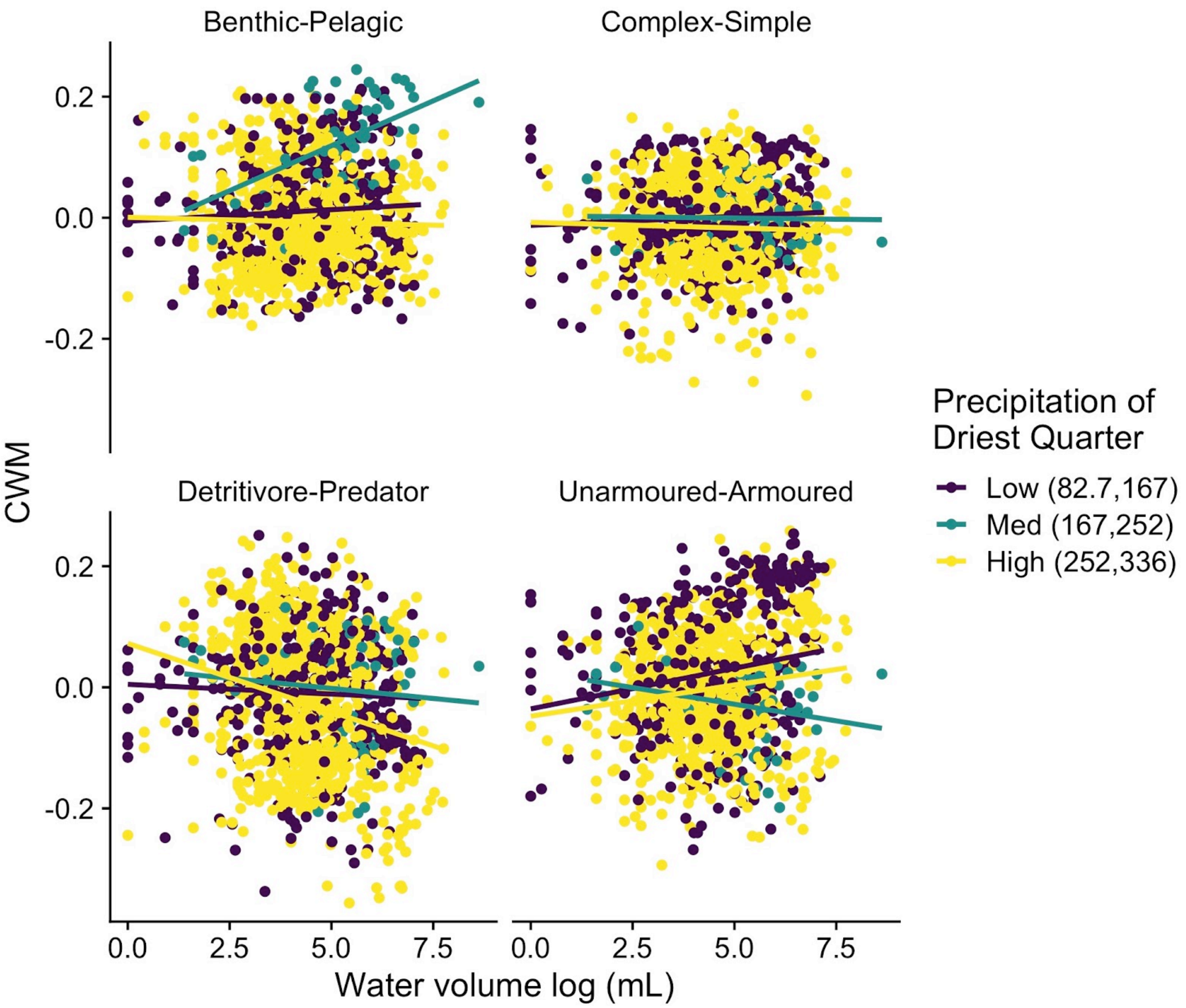


**Figure A11:** The four trait axes differ in their relationship with the total water volume which is also mediated by the precipitation in the driest quarter. Each point is the community weighted mean of a single bromeliad. The coloured lines are simple linear regressions intended only to improve the visualization of the data and not meant to be used for formal analysis since the CWM are multivariate.

**Table A3:** Synthetic trait composition (CWM) explained by local conditions in the fine-grained analysis and by climatic variables in the coarse-grained analysis. The analysis using the local conditions uses the CWM for each bromeliad. This analysis is blocked within each bioclimatic zone. The analysis using the biogeographic climatic variables uses the CWM for the species pool for each bioclimatic zone. This analysis used all the bromeliads for each field visit and did not sub-sample bromeliads.

| *Fine-grained analysis* | | |
|---|---|---|
| **Term** | **SS** | **P-value** |
| Total detritus | 0.213 | 0.812 |
| Canopy cover | 0.203 | 0.940 |
| Actual water | 0.123 | 0.974 |
| Residual | 39.838 | |
| Total | 40.373 | |
| Coarse-grained analysis | | |
| **Term** | **SS** | **P-value** |
| Mean Diurnal Range | 0.071 | 0.074 |
| Temperature Seasonality | 0.0481 | 0.116 |
| Precipitation of Driest | 0.041 | 0.320 |

| | | |
|---|---|---|
| Quarter | | |
| Precipitation Seasonality | 0.043 | 0.204 |
| Residual | 0.275 | |
| Total | 0.425 | |

**Table A4:** Synthetic trait composition explained by local conditions, biogeographic climatic variables and their interactions. This analysis used the CWM for each bromeliad. We did not include the interaction between canopy cover and climatic conditions because few bioclimatic zones had both open and closed canopy, therefore canopy cover would be confounded with bioclimatic zone. This analysis used all the bromeliads for each field visit and did not sub-sample bromeliads.

| *Local conditions* | | |
|---|---|---|
| **Term** | **SS** | **P-value** |
| Total detritus | 0.193 | 0.678 |
| Canopy cover | 0.219 | 0.824 |
| Actual water | 0.123 | 0.928 |
| *Climatic* conditions | | |
| Mean Diurnal Range | 0.095 | 0.148 |

| | | |
|---|---|---|
| Temperature Seasonality | 1.015 | 0.266 |
| Precipitation of Driest Quarter | 0.020 | 0.008 |
| Precipitation Seasonality | 0.862 | 0.312 |
| *Interactions between local conditions and climatic variables* | | |
| Total detritus: Mean Diurnal Range | 0.227 | 0.278 |
| Total detritus: Temperature Seasonality | 0.058 | 0.292 |
| Total detritus: Precipitation of Driest Quarter | 0.056 | 0.324 |
| Total detritus: Precipitation Seasonality | 0.152 | 0.922 |
| Actual water: Mean Diurnal Range | 0.430 | 0.184 |
| Actual water: Temperature Seasonality | 0.175 | 0.496 |
| Actual water: Precipitation of Driest Quarter | 0.409 | 0.068 |
| Actual water: Precipitation Seasonality | 0.071 | 0.492 |
| Residual | 36.266 | |
| Total | 40.373 | |
